# Subchondral H-type Blood Vessel Formation Aggravates Articular Cartilage Degeneration through LEP-LEPR Axis

**DOI:** 10.64898/2026.01.05.697672

**Authors:** Chao Yu, Jun Qin, Xi Zhao, Shitong Luo, Hao Zeng, Bo Xu, Jingjing Li

## Abstract

**Background:** Osteoarthritis (OA) is a chronic degenerative joint disease characterized by cartilage deterioration that has resulted in severe physical and economic costs for society. Risk factors for osteoarthritis encompass genetic predisposition, mechanical influences, obesity, inflammation, and metabolic problems. Recent reports indicate that the morphological alterations of subchondral bone evolve dynamically with the advancement of osteoarthritis. A significant alteration in subchondral bone is the development of aberrant blood vessels, particularly H-type vessels. H-type arteries are essential for preserving subchondral bone homeostasis. They support nutrients and oxygen to bone tissue and eliminate waste, thereby supporting the metabolic activities of osteoblasts and osteoclasts. However, the upstream factors governing H-type arteries are yet unknown.

**Methods:** The whole transcriptome analysis was performed by informatics which is further verified in clinical samples. Using LEP and LEPR knockout mice, the correlation between abnormal subchondral bone metabolism and LEP, LEPR, and H-type vessel formation were uncovered.

**Conclusions:** This study reveals that the lipid metabolism pathway was highly enriched on the OA lesion, and LEP and LEPR were markedly underexpressed in the subchondral bone on the side with severe OA lesions. The causal relationship of the LEP-LEPR-CD31 signalling pathway in aging and osteoarthritis models is revealed at the overall and molecular levels. This project will provide a new theoretical basis for the pathogenesis of early OA and new ideas and targets for the clinical treatment of OA by exploring the LEP-LEPR-CD31 signalling in osteoarthritis.

## INTRODUCTION

Osteoarthritis (OA) is a chronic, degenerative joint disorder marked by the deterioration of cartilage. It commonly manifests in weight-bearing joints and those subjected to heavy activity, serving as the primary cause of physical disability^1^. As human life expectancy rises, the global health and economic ramifications of osteoarthritis will persistently escalate in the future^2^. The pathological features of osteoarthritis include cartilage degradation, synovial inflammation, osteophyte development, and subchondral bone remodelling^3^. Notwithstanding the presence of established risk factors such as genetic predisposition, mechanical influences, obesity, inflammation, and metabolic disorders, the precise pathophysiology remains ambiguous, leading to suboptimal treatment success in most patients. Consequently, investigating the fundamental etiology of osteoarthritis has emerged as a critical subject requiring immediate attention^4^.

Recent evidence indicates that articular cartilage degradation is associated with aberrant subchondral bone remodelling^5–7^. Subchondral bone offers mechanical support to articular cartilage and collaborates with it to convey pressure loads, ensure joint congruence, and avert stress concentration inside the joint. In contrast to the slower remodelling rate of articular cartilage, subchondral bone exhibits a more rapid remodelling process in reaction to alterations in the mechanical environment^8^. The morphological alterations of subchondral bone evolve dynamically with the advancement of osteoarthritis. In the initial or advanced phases of osteoarthritis, subchondral bone remodelling is heightened, and the sites of bone rebuilding proliferate, resulting in a thinner and more porous subchondral bone plate. As the disease advances, the thickness of subchondral bone trabeculae diminishes, while the incidence of fractures and separations escalates. Active bone repair diminishes the thickness of the subchondral bone plate. In the advanced stage of osteoarthritis, bone remodelling diminishes, bone resorption declines, but bone synthesis increases comparatively. The subchondral bone plate and trabeculae thicken, the calcium-collagen ratio becomes imbalanced, and subchondral bone sclerosis escalates. The uncalcified cartilage exhibits gradual deterioration at this period. The coupling process between normal bone resorption and creation is disrupted in the advanced stage of osteoarthritis, resulting in a greater propensity for bone growth^9,10^. The mechanism that triggers aberrant subchondral bone production and the correlation between heightened subchondral bone remodelling and articular cartilage degradation in osteoarthritis remain ambiguous.

A significant alteration in osteoarthritis subchondral bone is the development of aberrant blood vessels, particularly H-type blood vessels^9^. H-type blood arteries possess distinctive morphology, positioning, and functionality. They are predominantly located in subchondral bone and are designated based on their H-type morphology. Their endothelial cells have elevated expression of platelet endothelial cell adhesion molecule-1 (PECAM-1/CD31) and neomycin (EMCN). H-type arteries are essential for sustaining subchondral bone homeostasis. They supply nutrients and oxygen to bone tissue and eliminate waste, therefore facilitating the metabolic functions of osteoblasts and osteoclasts^11^. Besides their function in bone metabolism, H-type arteries are linked to the pathogenesis of osteoarthritis (OA)^12^. Excessive vascularization, especially H-type arteries, in subchondral bone is deemed responsible for aberrant subchondral bone remodelling and cartilage deterioration in osteoarthritis. Additional research has demonstrated that alterations in H-type vessel density and morphology correlate with the onset and advancement of OA. The existence of subchondral H-type vessels correlates with elevated bone remodelling and enhanced subchondral bone permeability. This heightened permeability may facilitate the infiltration of inflammatory cells and exacerbate the pathogenesis of osteoarthritis^13,14^. Other investigations indicate that H-type arteries are considerably more prevalent in subchondral bone samples from patients with advanced osteoarthritis compared to those with mild or no osteoarthritis. The development of H-type vasculature in subchondral bone correlates with heightened expression of angiogenic factors and inflammatory cytokines, potentially exacerbating the progression of osteoarthritis^12,15,16^.

Leptin (LEP) is a hormone which is crucial for the regulation of energy equilibrium, hunger, and body mass. Adipose tissue produces and releases it, influencing many tissues, including bones, and positively affecting bone metabolism^17^. The leptin receptor (LEPR) is a membrane protein expressed in multiple tissues, including bone, and is crucial for modulating the effects of LEP ^17^. Upon binding to LEPR, LEP stimulates the signalling system that governs bone metabolism, enhances the proliferation and differentiation of osteoblasts, and suppresses osteoblast death, thereby augmenting bone production^18^. Furthermore, LEP decreases the development and activity of osteoclasts, hence diminishing bone resorption^19^. Deficiency in LEP or LEPR can result in alterations in bone metabolism and heightened bone loss, signifying that their modulation is crucial for sustaining bone health^20^. Furthermore, LEP and LEPR participate in the regulation of lipid metabolism. Deficiencies in LEP and LEPR might result in lipid metabolism abnormalities, augmented adipose tissue bulk, and elevated levels of free fatty acids (FFAs), potentially causing excessive adipocyte growth and chronic inflammation in adipose tissue^21^. Elevated concentrations of free fatty acids (FFAs) might catalyze the angiogenesis process by enhancing the expression of vascular endothelial growth factor (VEGF), so facilitating the proliferation and differentiation of endothelial cells^22,23^. Obesity and dysregulated fatty acid metabolism are significant risk factors for osteoarthritis. Will the LEP-LEPR axis influence the development of subchondral H-type blood vessels by modulating the concentration of FFAs in subchondral bone?

This study initially established the association between defective subchondral bone metabolism and LEP, LEPR, and H-type blood vessel development, based on LEP and LEPR knockout mice. The causal link of the LEP-LEPR-CD31 signalling pathway in aging and osteoarthritis models is elucidated. Our goal aims to establish a novel theoretical framework for the pathophysiology of early osteoarthritis and to propose innovative concepts and targets for its therapeutic therapy by investigating the LEP-LEPR-CD31 signalling pathway in osteoarthritis.

In this manuscript writing, we followed the guideline suggest by Transparency In The reporting of Artificial INtelligence – the TITAN guideline ^24^.

## RESULTS

### 1. LEP and LEPR are associated with subchondral bone pathological changes in OA

Clinically, patients with medial osteoarthritis classified grade 3-4 as Kellgren-Lawrence demonstrate considerable deterioration of the medial tibial plateau cartilage and subchondral bone sclerosis, while the lateral tibial plateau cartilage is predominantly intact, and the subchondral bone architecture is typically normal. To examine the relationship between the pathological characteristics of the medial subchondral bone and changes in the gene expression profile of the subchondral bone, we analysed the RNA sequencing data from GEO database (GSE51588), which includes the comprehensive genome expression profile chip data of the medial tibial plateaus (MT) and lateral tibial plateaus (LT) of the subchondral bone from 20 individuals with medial knee OA. We used the data from the LT of OA subchondral bone as the control group and MT as the experimental group, GSEA analysis was performed based on the recorded 23,002 gene expression data points. The results show that unlike the lateral subchondral bone, the fatty acid production, adipocytokine signalling pathway, and lipolysis regulation in the adipocytes of the medial subchondral bone were significantly inhibited (Figure 1A and 1C). The three pathways are all related to lipid metabolism, implying a hypothesis that the pathological changes in the medial subchondral bone are potentially connected to genes that regulate abnormal lipid metabolism.

**Figure 1:**
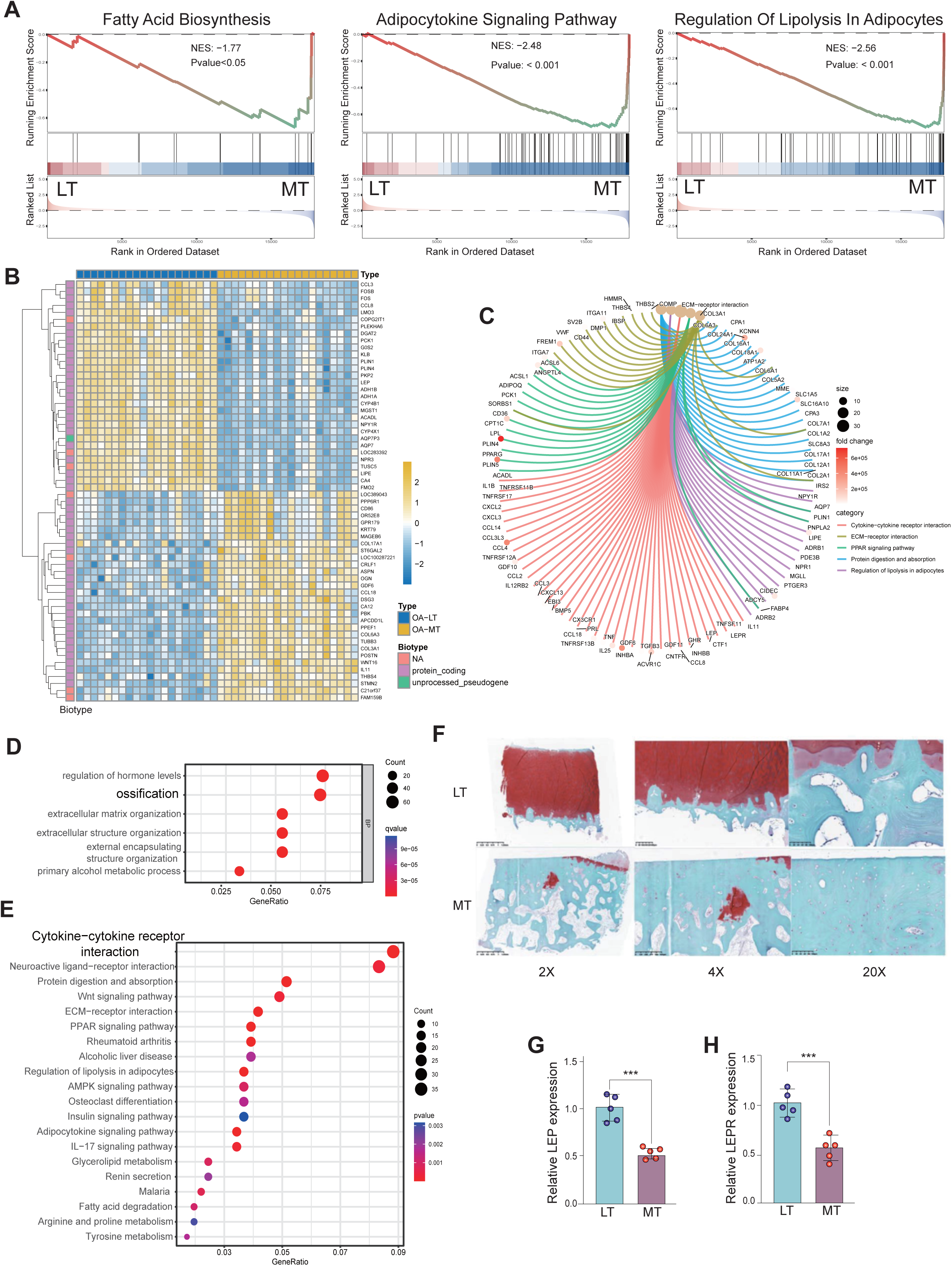
**A.** GSEA analysis of gene expression profile of medial subchondral bone (MT) and lateral subchondral bone (LT) in OA. **B**. Heat map of DEGs. **C**. Peacock map display of the important DEGs genes in the KEGG pathway. **D**. GO analysis of DEGs. **E**. KEGG pathway enrichment analysis of DEGs. **F**. Representative safranin O/fast green stain in samples from MT and LT subchondral bone of OA patients. **G and H**, PCR detection of the expression of LEP and LEPR in MT and LT subchondral bone of OA patients. * indicates P value<0.05, ** indicates P value<0.01, *** indicates P value<0.001.

To further explore the relationship between subchondral bone pathological changes and gene regulation, we used the GEO2R preprocessing core package of the GEO database to screen 23,002 genes in the dataset for differentially expressed genes (P<0.05) and obtained a total of 11,181 differentially expressed genes (DEGs) (Supplementary Figure 1A).

Then, we performed Gene ontology (GO) analysis and Kyoto Encyclopaedia of Genes and Genomes (KEGG) analysis based on DEGs (Figure 1D). Enriched biological process (BP) analysis shows that regulation of hormone levels and ossification terms are significantly enriched (Figure 1E). Besides, molecular function (MF) analysis suggests that receptor-ligand activity and signalling receptor activator activity were significantly enriched, collagen-containing extracellular matrix on cell components (CC) was significantly enriched, and cytokine-cytokine receptor interaction in KEGG pathway was significantly enriched. These results imply us to further narrow down the scope of key molecules.

To find the key genes regulating subchondral bone lesions, we further screened the genes with expression differences of more than 2 times and converted the fold change (FC) values by taking the logarithm (log2) to further narrow the range of differentially expressed genes. We obtained 1005 significantly differentially expressed genes (P<0.05, log21Z1 or 1Z -1), of which 422 genes were up-regulated, and 583 genes were down-regulated. The significantly differentially expressed genes were then constructed using the String protein database (https://cn.string-db.org) to construct a PPI network, and hub gene screening was performed using Cytoscape, obtaining 44 hubs such as LIPE, FABP4, LPL, LEP, and LEPR. Gene (Supplementary Figure 1B to 1D), combined with the results of GSEA, GO, and KEGG analysis, and combined with the heat map significantly different gene list (Figure 1B), it was found that LEP, LEPR molecules and signalling pathways can simultaneously meet the results of the above bioinformatics analysis. Therefore, we assume that LEP and LEPR may be related to abnormal lipid metabolism and pathological changes in subchondral bone.

To verify the results of bioinformatics analysis, we selected 5 human medial OA specimens (K-L grade 3-4) collected in the early stage. Through scanning observation after Safranin O-Fast Green staining, compared with the lateral tibial plateau, the cartilage defect on the medial lesion side was serious and the defect area was large. Subchondral bone sclerosis was seen, and many cell cavities were seen inside (Figure 1F). At the same time, we extracted RNA from the medial and lateral subchondral bone tissues, respectively, and performed PCR detection of LEP and LEPR genes. Consistently, it was found that the expression levels of LEP and LEPR in the medial subchondral bone decreased (Figure 1G and 1H).

As for this, we verified the gene expression changes of LEP and LEPR through clinical samples, which were consistent with the trend of the previous bioinformatics analysis results, suggesting that the decrease in LEP and LEPR expression levels may be related to pathological changes such as subchondral bone sclerosis on the lesion side and OA.

### 2. Early OA pathological changes in subchondral bone of mice with LEP and LEPR gene deletion

To better understand the function of LEP and LEPR genes in subchondral bone, we generated 6–7-week-old wild type mice (db/db background mice, as control group), db/db mice (LEPR homozygous mutation), and ob/ob mice (LEP homozygous mutation). Compared with wild type mice, db/db and ob/ob mice showed strong appetite and slow movements. Grossly, db/db and ob/ob mice were obese and had subcutaneous fat accumulation (Figure 2A). Although no obvious joint deformities or gait abnormalities were observed either in db/db or ob/ob mice (Figure 2B), the bone volume, tissue volume, bone volume fracture and bone surface area are all decreased in the mice with LEP/LEPR homozygous mutation (Figure 2C). Micro CT scanning shows decreased number of trabecular bones in db/db and ob/ob mice Figure 2D and 2E)^25,26^.

**Figure 2:**
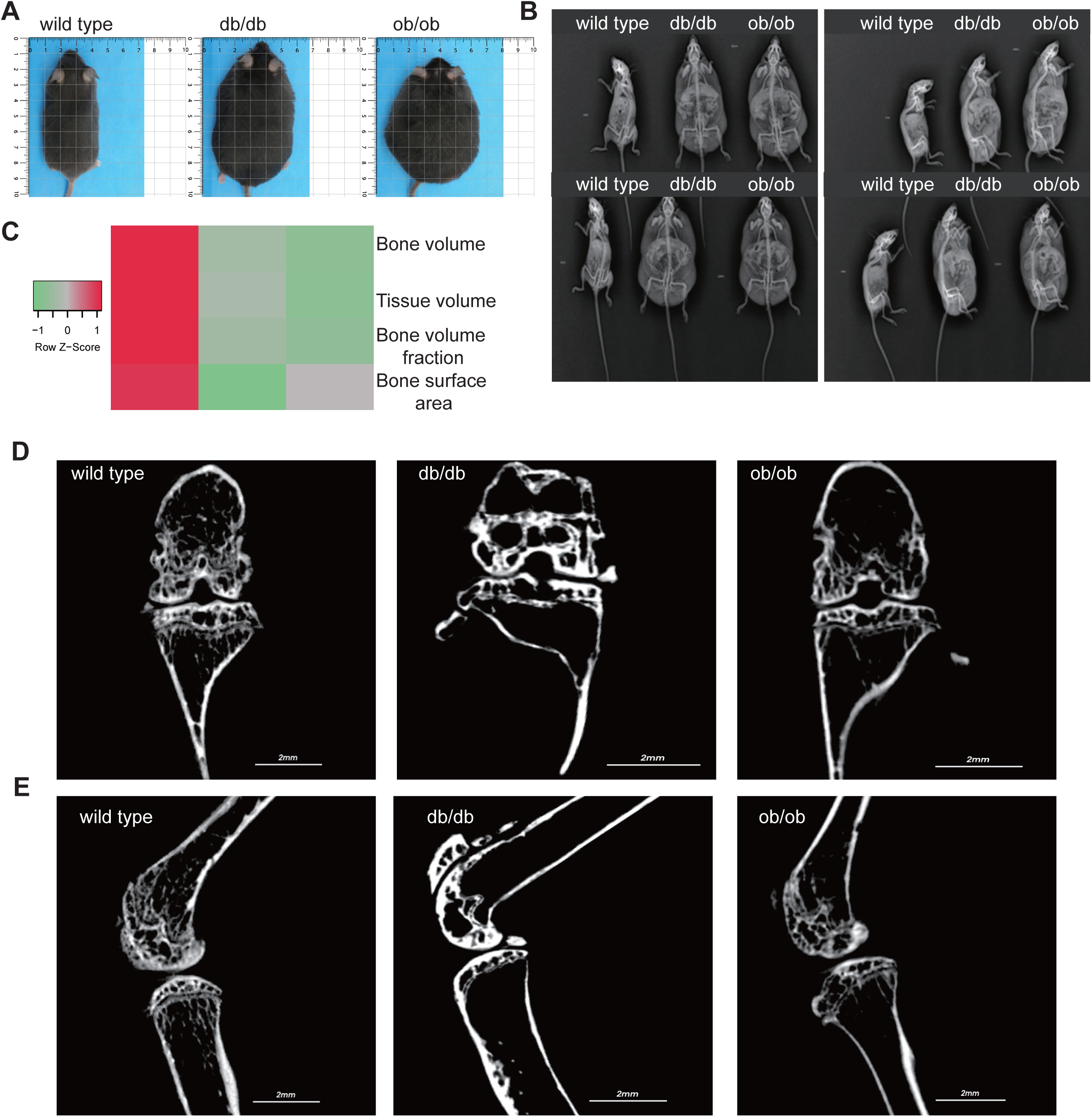
**A**. General appearance of 6-month-old wild type, db/db, ob/ob mice. **B**. Front and side X-ray display of bones and joints of 6-month-old wild type, db/db, ob/ob mice. **C**. Heatmap showing virous parameter of micro-CT scanning in the transgenic mice. **D**. Representative bone tissue micro-CT scan image from the front view. **E**. Representative bone tissue micro-CT scan image from the side.

### 3. Either LEP or LEPR is essential for joint microstructure maintaining

To ascertain the impact of LEP and LEPR loss on joint microstructure, we fixed, decalcified, and embedded the joints of the sacrificed mice, obtained sagittal sections, and examined the morphology of the articular cartilage using safranin O-fast green staining. Upon examination, the joint surfaces of the three mice exhibited a smooth texture, devoid of fissures or indications of deterioration. Safranin staining revealed varying degrees of loss across each group; however, no significant differences were seen (Figure 3A). We observed a notable increase in fat vacuoles and a reduction in bone mass in the subchondral bone of db/db and ob/ob mice, particularly in db/db mice. This resembles the occurrence of lipid accumulation, cystic alterations, and cavity formation in the subchondral bone on the affected side, frequently reported clinically; nevertheless, the data revealed no indication of harm to the cartilage structure in mice due to lipid accumulation. To enhance comprehension of the development of fat vacuoles beneath the cartilage resulting from the absence of LEP and LEPR genes, To further investigate the phenomena of collagen, decrease, we employed Sirius red staining. Collagen fibers in the tissue appeared red using a standard optical microscope; however, assessment of different collagen fibers was enhanced using a polarized light microscope. In accordance with the findings from Masson staining, lipid buildup was evident in the subchondral bone of db/db and ob/ob mice, and there was a marked reduction in the red collagen fibers within the cartilage and subchondral bone (Figure 3B). Collagen is a crucial component for preserving bone strength and supplying nutritional support to bones. Collagen can facilitate the deposition of calcium salts in bone tissue. db/db and ob/ob mouse models Sirius red and Masson staining reveal a reduction in subchondral bone collagen, potentially resulting in heightened bone brittleness and precipitating conditions such as osteoporosis and fractures.

**Figure 3.**
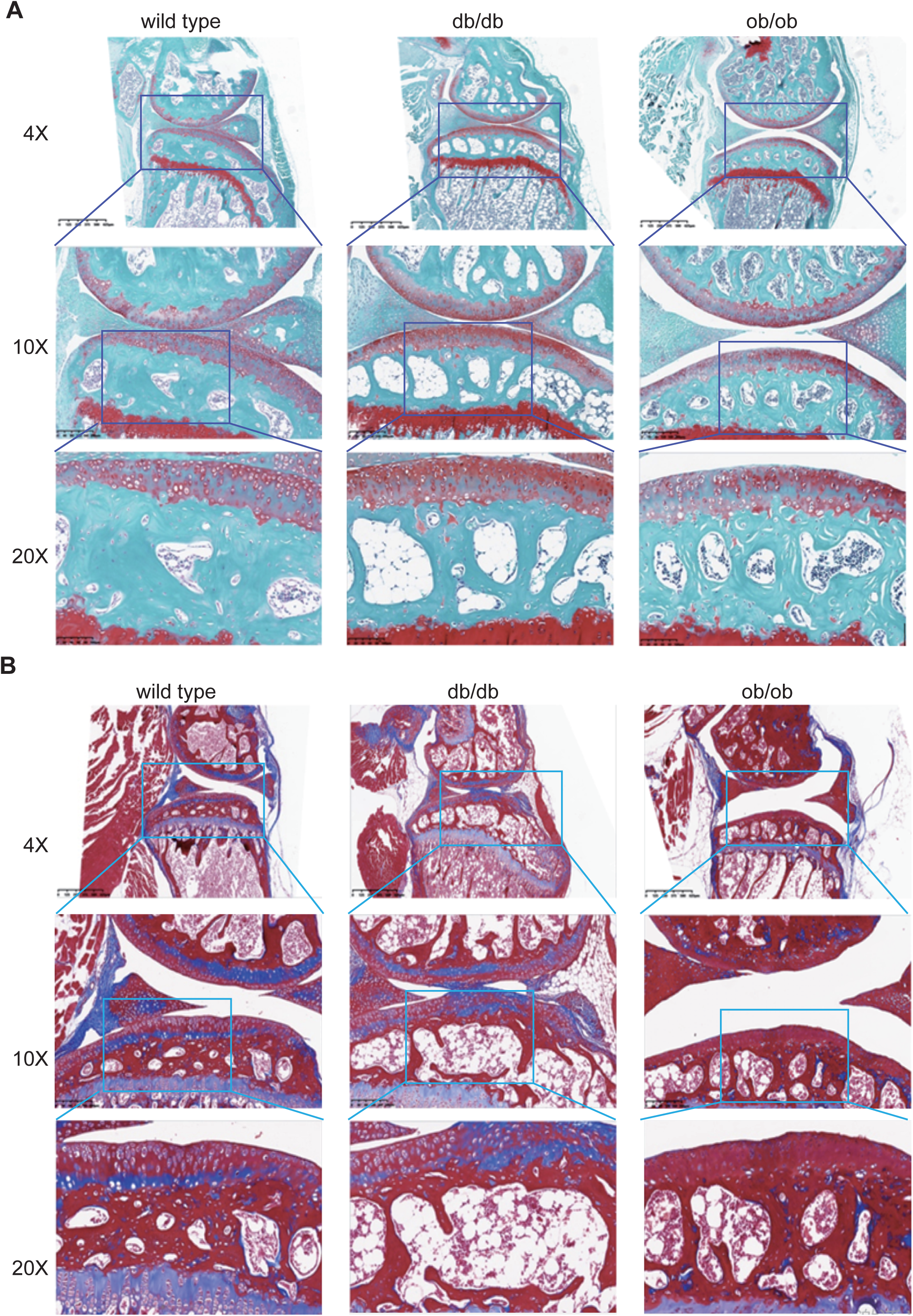
**A**. Representative safranine o/fast green stain of 6-month-old wild type, db/db, ob/ob mice joint. **B**. Representative Masson stain of 6-month-old wild type, db/db, ob/ob mice joint.

In both the early and severe stages of osteoarthritis, the activity of subchondral osteoclasts is elevated, resulting in active subchondral bone remodelling^27,28^. To assess the impact of postnatal loss of LEP and LEPR on osteoclasts in subchondral bone, we employed TRAP staining to examine the distribution and proportion of osteoclasts. Osteoclasts were stained crimson by TRAP and situated in the cytoplasm. The subchondral bone of db/db and ob/ob mice exhibits a limited presence of wine-red patches, however their distribution is not extensive. Nonetheless, a significant quantity of wine-red cells was observed in db/db and ob/ob animals beneath the tidal line (Figure 4A), signifying an increase in osteoclasts in this region and aggressive bone breakdown, which aligns with the characteristics of early osteoarthritis.

**Figure 4:**
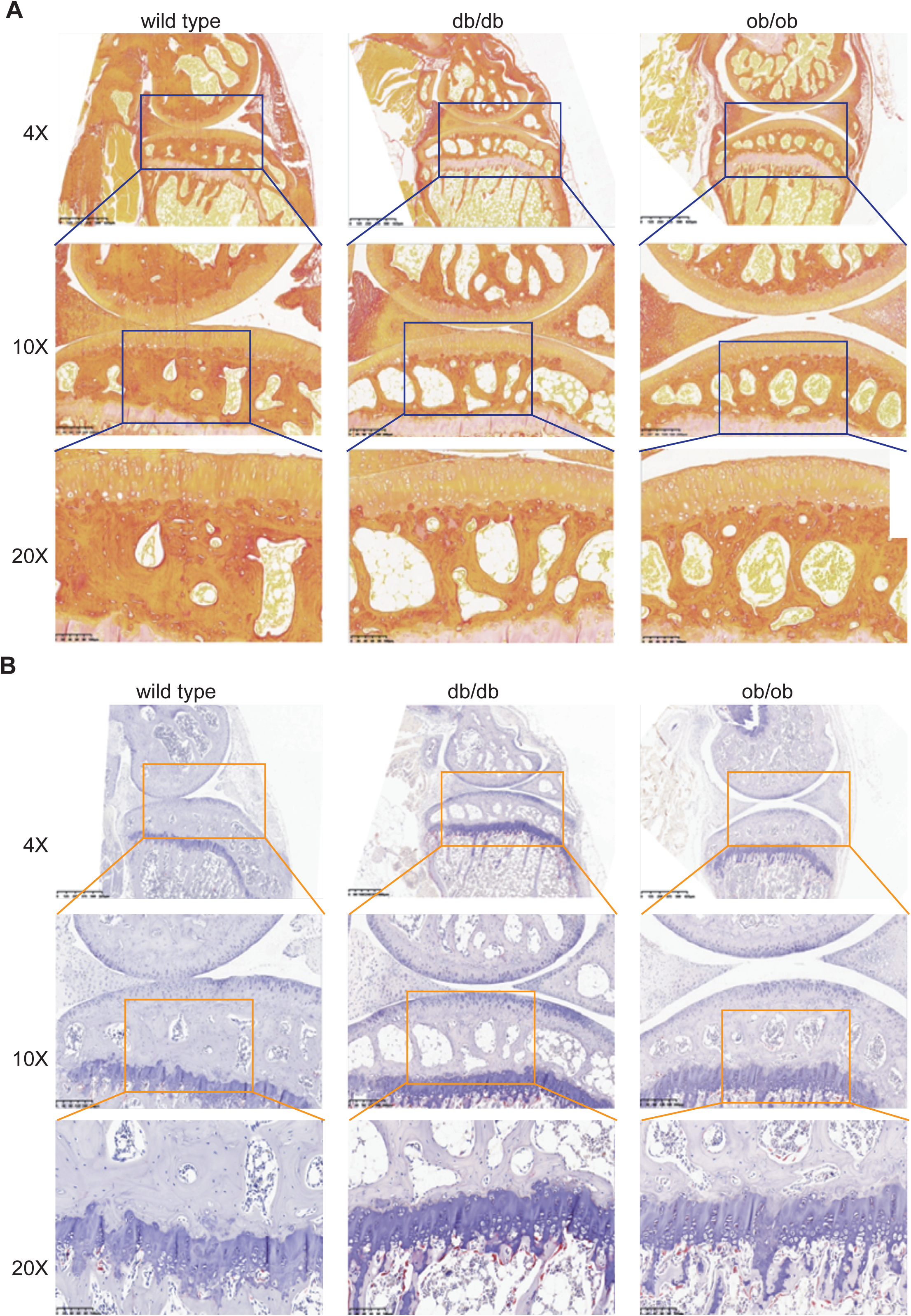
**A**. Representative Sirius red staining of 6-month-old wild type, db/db, ob/ob mice joint. **B**. Representative TRAP staining of 6-month-old wild type, db/db, ob/ob mice joint.

The histological staining revealed that the postnatal loss of LEP and LEPR resulted in lipid metabolic abnormalities in subchondral bone, leading to lipid buildup. While this occurrence did not alter the structure of cartilage in its normal form, subsequent analysis revealed a reduction in the amount of subchondral collagen fibres and increased osteoclast activity, aligning with the findings of early clinical osteoarthritis.

Deletion of the LEP and LEPR genes expedites cartilage degradation in murine models. Cartilage degeneration correlates with atypical subchondral bone remodelling. To ascertain if the alterations in the subchondral bone induced by LEP and LEPR gene deletion impact the chondrocytes in the upper cartilage, we employed toluidine blue to stain the mouse joints. Post-scan analysis revealed that the overall architecture of the cartilage and subchondral bone aligned with the findings from Safranin O-Fast Green staining, indicating an increase in vacuole count within the subchondral bone, while no significant cracks or defects were observed in the cartilage. Notably, the quantity of hypertrophic chondrocytes stained by toluidine blue in the cartilage of db/db and ob/ob mice exhibited a marked increase (Figure 4B).

Col-X is a molecular marker expressed during chondrocyte differentiation and hypertrophy, primarily involved in the transformation of chondrocytes into osteoid cells. In the progression of osteoarthritis, chondrocytes progressively deteriorate in both morphology and function, subsequently secreting excessive collagen type X, which exacerbates the structural and functional impairment of cartilage [24]. We employed immunohistochemical labeling to examine the expression of the chondrocyte-specific marker Col-X. In accordance with the findings from toluidine blue staining, the expression of Col-X in the cartilage of db/db and ob/ob mice was markedly elevated (Figure 5A).

**Figure 5:**
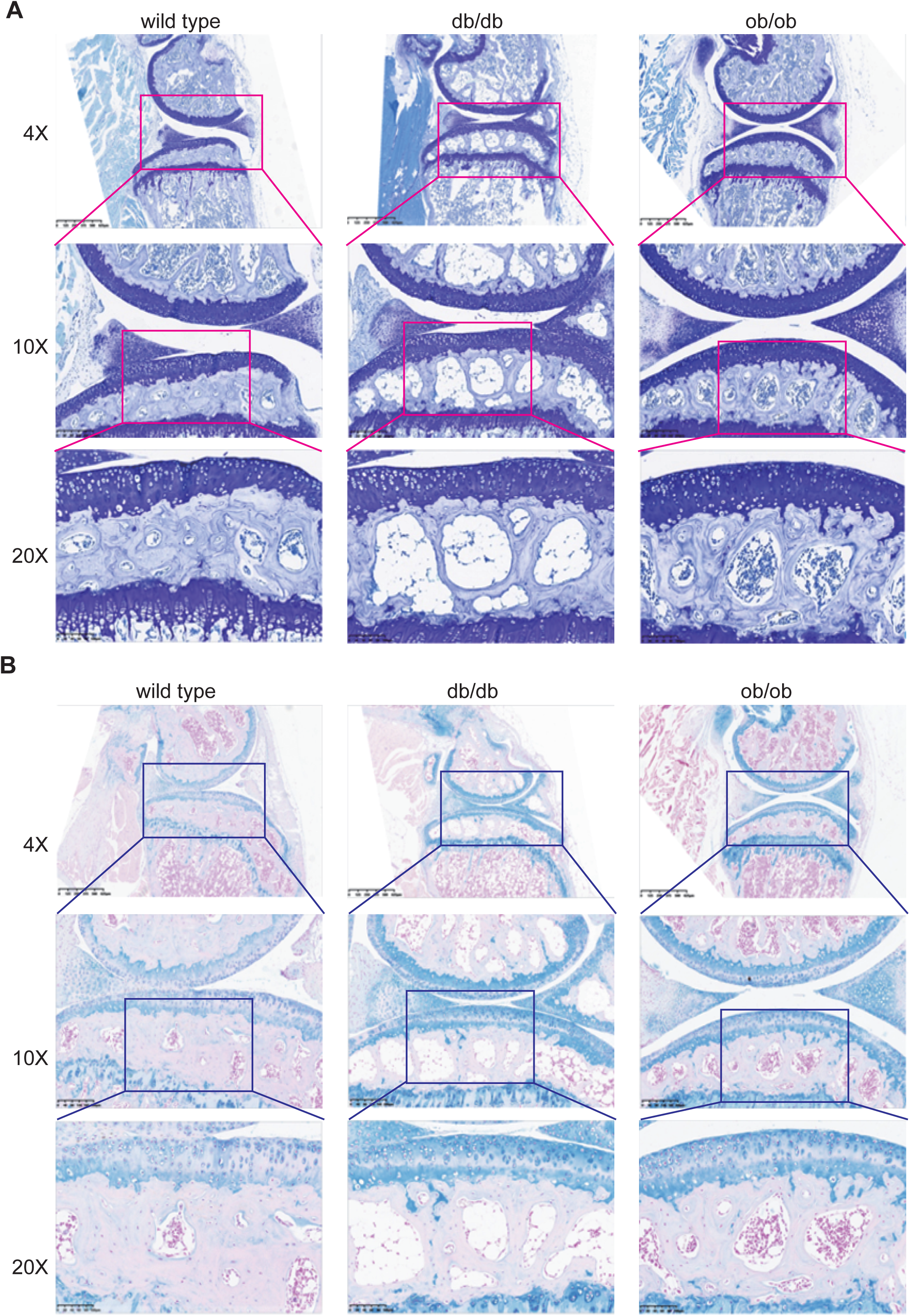
**A**. Representative toluidine blue staining of 6-month-old wild type, db/db, ob/ob mice joint. **B**. Representative Alcian Blue staining of 6-month-old wild type, db/db, ob/ob mice joint.

To elucidate the impact of chondrocyte hypertrophy resulting from the absence of LEP and LEPR on cartilage components, we employed Alcian blue (A-B) staining to examine alterations in cartilage tissue composition. Alcian blue is a specialized dye for acidic mucopolysaccharides. It causes acidic chondroitin, glycosaminoglycans, and mucopolysaccharides in the tissue to exhibit a blue hue, whereas neutral glycoproteins and collagen display a red hue, so differentiating acidic glycoproteins from neutral glycoproteins. In comparison to the cartilage of wild type mice, the blue coloration of the articular cartilage in db/db and ob/ob mice was markedly more pronounced (Figure 5B), suggesting a large increase in the concentration of acidic glycoproteins. Their anomalous elevation may result in cartilage degradation and exacerbation of the inflammatory response^29,30^. The findings indicate that the postnatal deficiency of LEP and LEPR results in lipid accumulation within the subchondral bone, an elevation in cartilage mast cells, an increase in intracartilage acidic glycoproteins and mucopolysaccharides, alterations in cartilage constituents, and initial indications of cartilage degeneration.

## 4 Elevated H-type blood vessels in the subchondral bone of mice with deletions of the LEP and LEPR genes

The density of H-type blood vessels in the subchondral bone correlates with the onset and progression of OA. Abnormal lipid metabolism has been linked to the growth of H-type blood vessels. To further examine whether lipid accumulation in the subchondral bone resulting from the loss of LEP and LEPR genes stimulates the formation of H-type blood vessels, we employed immunofluorescence (IF) and immunohistochemistry (IHC) to detect the angiogenesis markers VEGFR, HIF-1α protein, and the specific marker CD31 protein associated with H-type blood vessels. The elevation of VEGFR enhances the sensitivity of H-type vascular endothelial cells to VEGF signals^31^. The green fluorescence of VEGFR surrounding the subchondral bone in db/db mice exhibited a considerable rise, while the subchondral bone in ob/ob mice also shown an upward trend compared to wild type animals (Figure 6A). Hypoxia-inducible factor-1α (HIF-1α) is a transcription factor that is activated in response to hypoxic conditions. HIF-1α has been reported to induce the development of H-type blood arteries. We employed immunofluorescence to ascertain the expression level of HIF-1α in subchondral bone. In accordance with the VEGFR findings, robust green fluorescent protein expression was observed in the subchondral bone of db/db and ob/ob mice, particularly pronounced in the subchondral bone of db/db mice (Figure 6B).

**Figure 6:**
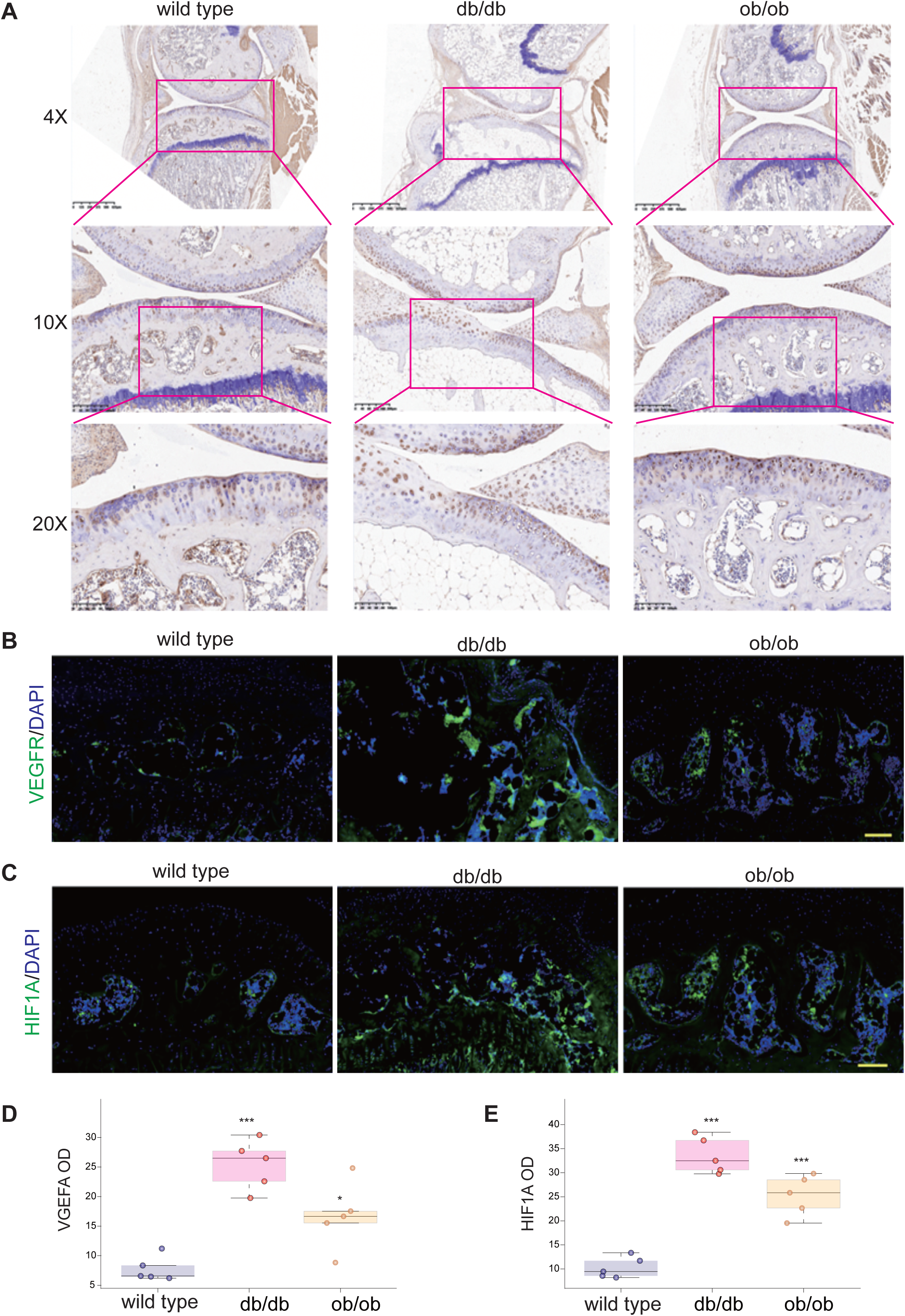
**A**. The expression of Col-X in the joints of 6-month-old wild type, db/db, ob/ob mice was detected by IHC. **B**. The expression of VEGFR in the joints of 6-month-old wild type, db/db, ob/ob mice was detected by immunofluorescence. **C**. The expression of HIF-1α in the joints of 6-month-old wild type, db/db, ob/ob mice was detected by immunofluorescence. D. Quantification analysis of VGEFA optical density. E. Quantification analysis of HIF1A optical density. * indicates P value<0.05, ** indicates P value<0.01, *** indicates P value<0.001.

CD31 protein serves as a unique marker for H-type vascular endothelial cells. We employed immunohistochemistry to stain the subchondral bone of mice (Figure 7A). In accordance with the distribution patterns of VEGFR and HIF-1α in the three mouse models, CD31 exhibited elevated expression in the subchondral bone of db/db and ob/ob animals(Figure 7B), indicating an over proliferation of H-type blood vessels in the subchondral bone of these mice.

**Figure 7:**
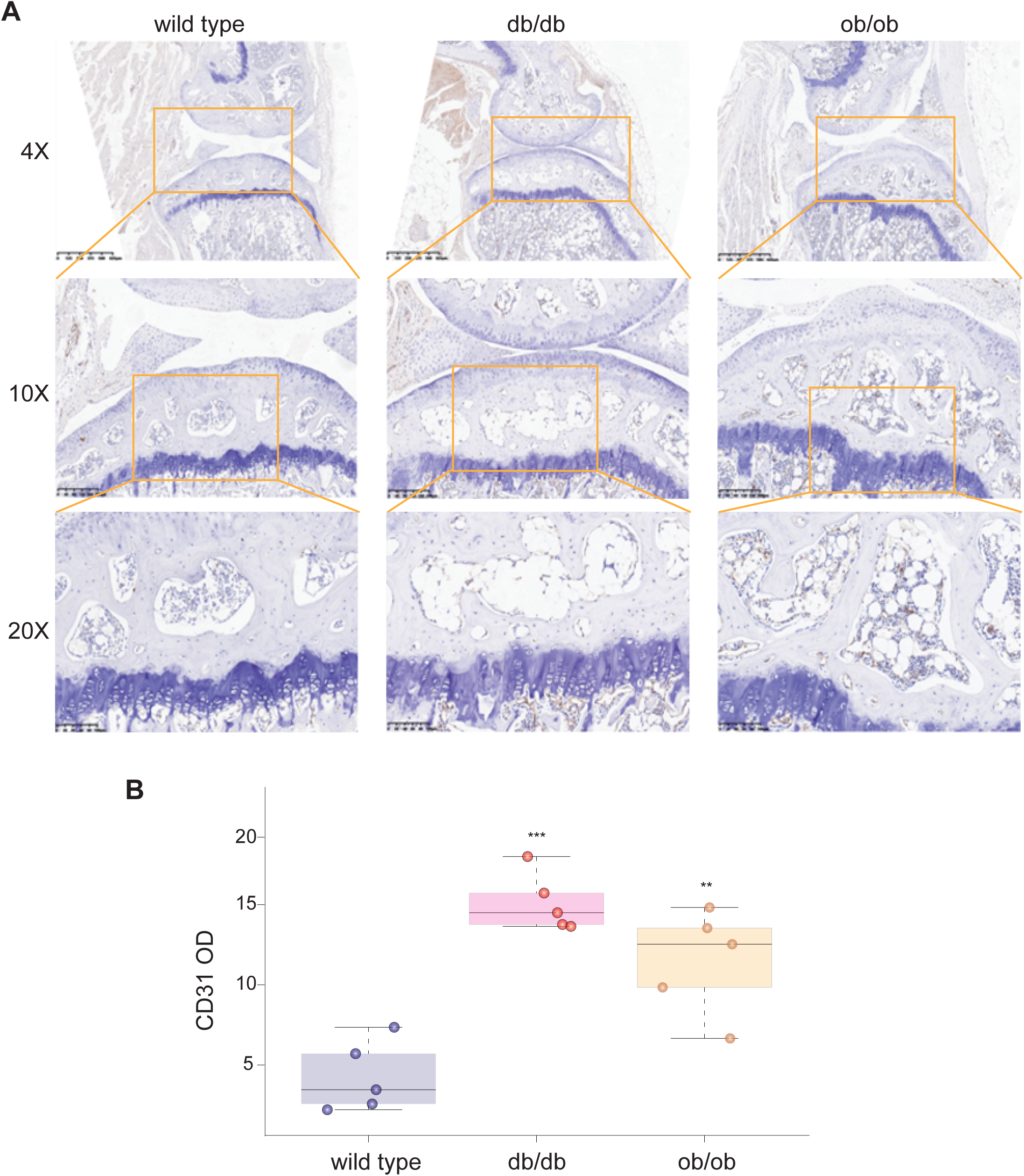
**A**. The expression of CD31 in the joints of wild type, db/db, ob/ob mice was detected by IHC. **B**. Quantification analysis of CD31 optical density. * indicates P value<0.05, ** indicates P value<0.01, *** indicates P value<0.001.

## DISSCUSION

In order to illustrate the upstream factors regulating H-type vessels, we developed a mouse model with postnatal deletion of LEP and LEPR to assess the impact of LEP and LEPR signalling on the bone architecture and collagen ratio of subchondral bone, as well as on chondrocyte hypertrophy and alterations in intracartilage glycoproteins, which align with the initial modifications in subchondral bone and cartilage observed in human osteoarthritis. Simultaneously, we confirmed that the influence of LEP and LEPR on subchondral bone may be mediated through the modulation of lipid metabolism, subsequently affecting H-type blood vessels, so offering valuable insights for the future understanding and prevention of OA.

Recent research increasingly indicates that aberrant lipid metabolism, particularly abnormalities of fatty acid metabolism, may contribute to the onset and progression of OA^32–34^. Aberrant lipid metabolism facilitates the onset and advancement of osteoarthritis by multiple pathways, including inflammation, cartilage degradation, subchondral bone sclerosis, and disturbances in chondrocyte anabolism and catabolism^35–37^. Lipids are essential constituents of cell membranes and serve as signalling molecules that modulate several cellular activities^38^. Aberrant lipid metabolism may result in lipid deposition in joints, causing inflammation and cartilage deterioration. Lipids, including free fatty acids and cholesterol, initiates cartilage deterioration by enhancing the synthesis of inflammatory cytokines and degradative enzymes^39^. Lipid metabolism influences the equilibrium between chondrocyte anabolic and catabolic activities. Dysregulation of lipid metabolism may also influence the activity of enzymes critical for collagen and proteoglycan production, which are vital for cartilage integrity^40^. Adipose tissue serves as the primary location for lipid metabolism and synthesizes various adipokines, including adiponectin, leptin, and resistin, which influence cartilage metabolism, inflammation, and angiogenesis, so facilitating the onset and advancement of osteoarthritis^41,42^. Lipid metabolism problems concurrently result in an imbalance between lipogenesis and lipolysis^43^. Lipids accumulate in the subchondral bone, potentially resulting in subchondral bone sclerosis, a hallmark of osteoarthritis^44^. This investigation revealed significant lipid buildup in the subchondral bone of db/db and ob/ob mice. The buildup of lipids diminished the amount of native subchondral bone tissue. Masson staining and Sirius red staining revealed a reduction in the collagen content of the subchondral bone. The reduction in collagen was correlated with a decline in bone mass, resulting in a thinner and more porous subchondral bone plate. It may exhibit increased susceptibility to fracture and separation under circumstances of trauma or severe external force. Concurrently, we also noted that the presence of active osteoclasts in the subchondral bone of ob/ob mice signifies that the subchondral bone undergoes active transformation and exhibits an increase in bone remodelling sites, resembling the characteristics of subchondral bone in both early and severe stages of osteoarthritis. Despite the absence of observable cartilage loss or destruction in mice with postnatal LEP and LEPR impairment, we noted an increase in mast cells and acidic glycoproteins inside the cartilage tissue, suggesting that their cartilage degeneration occurred at a more accelerated rate compared to normal mice. Consequently, it can be inferred that postnatal depletion of LEP and LEPR influences the distribution and trabecular architecture of subchondral bone, and these alterations in subchondral bone subsequently impact chondrocyte degeneration. This experiment did not explicitly determine the type of collagen in the subchondral bone that was diminished, nor the particular acidic glycoprotein in the cartilage that was reduced, together with the precise impacts on cartilage degeneration, necessitating future investigations for clarification.

Osteogenesis is crucial for preserving the integrity and functionality of the mammalian skeletal system. Osteogenic dysfunction and anomalies can result in numerous bone disorders^45,46^. A subtype of capillaries linked to osteogenesis has recently been identified as type H vasculature, characterized by elevated expression of CD31 and endomucin (Emcn)^47^. Type H vessels are situated adjacent to the metaphyseal growth plate and inside the periosteum and endothelium of the diaphysis. Type H arteries are closely encircled by osteoprogenitor cells that express the transcription factor Osterix, a powerful enhancer of bone production^48,49^. Type H vasculature actively facilitates bone formation by secreting substances that promote the proliferation and differentiation of osteoprogenitor cells^50,51^. Hypoxia-inducible factor 1α (HIF-1α), Notch, and VEGFR signalling are molecules significantly produced by type H vascular endothelial cells, which might enhance sensitivity to VEGF, hence promoting angiogenesis^47,52^. Angiogenesis in subchondral bone may significantly contribute to the etiology of osteoarthritis. The clusters of type H arteries surrounding subchondral bone islands in a murine model of osteoarthritis^53^. Type H vasculature is elevated in the subchondral bone of osteoarthritis model mice and elderly animals^54,55^. Activation of mTORC1 in cartilage enhances VEGF-A production in articular chondrocytes and facilitates the development of H-type blood vessels in subchondral bone, hence exacerbating osteoarthritis^56^. H-type angiogenesis in subchondral bone may be a significant characteristic of early osteoarthritis, and that the quantity of H-type blood vessels signify the extent of subchondral bone proliferation and remodelling^57,58^. In this work, we noted an elevation in the expression of CD31, a definitive marker of H-type blood vessels in the subchondral bone of mice with postnatal deletion of LEP and LEPR. This was accompanied by heightened expression and distribution of VEGFR and HIF-1α, suggesting an increase in the proliferation of H-type blood vessels in the subchondral bone of these mice. This may result from the heightened destructive impact of osteoclasts on subchondral bone, leading to an elevation in the production of substances that promote the formation of H-type blood vessels and augmented bone remodelling. Furthermore, it may also result from direct stimulation following the deletion of LEP and LEPR, or from the growth of H-type blood vessels induced by lipid metabolic problems. The precise mechanism requires additional investigation.

Trauma is one of significant risk factors in the onset of osteoarthritis^59^. Articular cartilage exhibiting deterioration demonstrates a sluggish or imbalanced reparative process following damage. This experiment previously demonstrated that mice with postnatal LEP and LEPR impairment exhibited weight growth, increased adiposity, H-type angiogenesis in subchondral bone, and cartilage degradation, all of which are risk factors for OA. Following cartilage damage, malfunction in cartilage repair or aberrant reconstruction of subchondral bone may arise due to an increase in H-type blood vessels in the subchondral bone, exacerbating cartilage degradation. Nonetheless, following the simulation of DMM surgery in the mice, we did not see significant OA cartilage degradation, which contradicted our anticipated outcomes. This investigation is warranted due to the evident obesity in db/db and ob/ob mice, particularly characterized by significant abdominal fat buildup. In a static condition, the interaction between abdominal adipose tissue and the base of the cage alleviates joint stress, rendering the DMM model insignificant in the cartilage-to-cartilage contact; conversely, db/db and ob/ob mice may exhibit reduced physical activity and remain in a static state for extended periods, resulting in a lack of relative mobility among joints. Consequently, we did not include images of mice post-DMM surgery in this study.

In conclusion, experimental confirmation of mice with LEP and LEPR deficiencies confirmed the link between the LEP-LEPR axis and subchondral bone H-type angiogenesis as well as articular cartilage degradation. The LEP-LEPR axis may regulate H-type angiogenesis and exacerbate articular cartilage degradation.

## Methods and materials

### 1. Animal assay

Experimental mice including wild-type (WILD TYPE-Leprem2Cd479/Gpt), diabetic (db/db) (WILD TYPE-Leprem2Cd479/Gpt), and obese (ob/ob) (B6/JGpt-Lepem1Cd25/Gp) mice, aged 6-7 weeks, were acquired from Jiangsu Jicui Yaokang Biotechnology Co., Ltd. and maintained in the SPF-level laboratory of the Experimental Animal Center at Chongqing Medical University. The mice were maintained at a constant temperature of 22±2IZ, relative humidity of 55±5%, and a light-dark cycle of 12 hours. Throughout the breeding procedure, mice were ensured unrestricted access to water and food, and all activities were conducted in compliance with the protocol sanctioned by the Animal Ethics and Use Committee of Chongqing Medical University (No.2021.5.28/LL-202133). This work has been reported in accordance with the ARRIVE guidelines (Animals in Research: Reporting In Vivo Experiments)^60^.

### 2. Data source and preprocessing

The GSE51588 gene expression profile dataset was obtained from the GEO database, comprising 40 subchondral bone samples from OA tibial plateaus (20 lateral vs 20 medial). The annotation data on the platform indicates that the probes were transformed into their respective gene symbols. The dataset was standardized using R (version 4.1.3, www.R-project.org) and the core preprocessing module in GEO2R. Subsequent GSEA, GO, and KEGG analyses were conducted using R packages, with P values < 0.05 being statistically significant.

### 3. Specimen Processing

Human sample collection was approved by the Ethics and Committee of Chongqing Medical University (2023-021). Human tissue specimens and murine knee joints were fixed in 4% paraformaldehyde for 36-48 hours, decalcified using 20% ethylenediaminetetraacetic acid (EDTA with pH 7.4) for 21 days, embedded in paraffin, and sagittal slices of the murine joints were prepared at a thickness of 7μm. The samples were stained with Safranin O-fast green, Toluidine blue, Sirius red, Masson, TRAP, and Alcian blue, among others, and subjected to immunohistochemistry and immunofluorescence techniques.

### 4. Toluidine blue staining

Slides were produced as described above, thereafter undergoing dewaxing and hydration. The slides were thereafter submerged in water briefly and incubated in a 0.25% toluidine blue dye solution for 2 to 5 minutes. Subsequently, excess dye solution was eliminated with water; the slices were rinsed with water and subsequently examined under a microscope using 0.1% glacial acetic acid. Cartilage and mast cells have distinct coloration, whereas the backdrop transitions to a light blue tint. Rinsing with tap water to cease differentiation, followed by drying with air. The slides were further dehydrated with anhydrous ethanol, rendered translucent with xylene, and then encapsulated in neutral gum. Toluidine blue can provide a purple-red coloration to chondroitin sulfate in cartilage tissue. This technique is utilized to investigate abnormal changes and the distribution of mast cells, subsequently analyzing the morphological structure of cartilage.

### 4. TRAP staining

A TRAP working solution was formulated by sequentially incorporating 50 μL of para-fuchsin solution (G1050-2), 50 μL of sodium nitrite solution (G1050-3), 100 μL of AS-BI phosphate substrate solution (G1050-4), and 1.8 mL of reaction buffer (G1050-1) into a centrifuge tube. Meticulously combine at each phase, thereafter filter using a 0.45μm needle filter membrane, followed by dewaxing the paraffin sections and rinsing with distilled water for several minutes. The slides were subsequently outlined using a histochemical pen and positioned in a humidified chamber. An appropriate volume of distilled water was added and incubated at 37°C for 2 hours. After the removal of water, the pre-prepared TRAP working solution was enough applied to cover the slides and incubated at 37°C in darkness for 20-30 minutes. Subsequently, the slides underwent the standard dehydration and mounting procedures. TRAP staining in osteoclasts displays a wine-red hue and is located within the cytoplasm.

### 5. Masson staining

In accordance with standard protocol, dewax the paraffin sections, drain them, and immerse them in solution A overnight; Following a 30-minute incubation at 65°C, rinse the sections with tap water the subsequent day to eliminate any residual solution A; Equal volumes of solutions B and C must be amalgamated, followed by a one-minute staining period and a subsequent fifteen-second washing with water. Differentiate for one to two minutes using a 1% hydrochloric acid alcohol solution (concentrated hydrochloric acid: anhydrous ethanol in a 1:100 ratio). Note: that the cell nucleus appears gray-black, but the backdrop is nearly colorless or light gray. Rinse with water and expel excess moisture to prevent differentiation. The specimen must be immersed in solution D for 6 to 8 minutes, ceasing when the tissue exhibits a vivid red coloration. Rinse with water to eliminate any excess liquid, then immerse in solution E for approximately 1–2 minutes. Note that while the other fibers are red, the collagen fibers exhibit a mild red hue. Rinse three times for approximately eight seconds each with 1% glacial acetic acid after immersion in F solution for two to thirty seconds. Employ anhydrous ethanol for dehydration for approximately five, ten, and thirty seconds. After 30 minutes and 2 minutes of n-butanol dehydration, clarify the slide with xylene and affix it using neutral gum. Following Masson’s trichrome staining, collagen fibers exhibit a spectrum from sky blue to deep blue, red blood cells appear light red, while cellulose, muscle fibers, cytoplasm, and keratin display hues ranging from red to purplish red.

### 6. Sirius Red Staining

Slides were produced as described above. Slides were immersed in Sirius red complex staining solution for 5-10 minutes. Then rinsing the slides with water, dehydrating using n-butanol for 30 seconds and 2 minutes, rendering transparent with xylene, and mounting the slide with neutral glue.

Collagen fibers in tissues are dyed red under a standard optical microscope. Observation using a polarized light microscope helps differentiate among distinct fiber kinds.

### 7. Alcian blue staining

Slides were produced as described above. The slides were immersed in Alcian blue dye solution for 15-30 minutes and rinsed with tap water. Submerge parts in nuclear quick red dye solution for 3 minutes and rinse with tap water. Dehydrate sections using anhydrous ethanol at three intervals of five minutes, clarify with xylene for two intervals of five minutes, and mount with neutral gum. Results Interpretation: Alcian blue is a particular stain for acidic mucins, rendering acidic mucopolysaccharides in the cytoplasm blue and neutral glycoproteins red, thus differentiating acidic glycoproteins from neutral glycoproteins.

## Acknowledgements

The report was supported by Chongqing Natural Science Foundation (CSTB2023NSCQ-MSX0640) and Chongqing Municipal Health Commission (2025MSXM081). Jingjing Li is supported by the National Natural Science Foundation of China (Grant No. 82104289), Shandong Provincial Health Commission(M-2022053), Science and Technology Innovation Plan from Weifang Medical University (041004), Yuandu Scholar Grant of Weifang City to LJJ, Weifang Science and Technology Bureau Plan Project (2021YX081), Science and technology project jointly established by the Science and Technology Department of the State Administration of Traditional Chinese Medicine (GZY-KJS-SD-2023-079), Shandong Provincial Medical Association Young Talent Promotion Project (2023_GJ_0039). Science and Technology Innovation Plan from Weifang Medical University (041011).

## Author’s contributions

Jingjing Li and Chao Yu conceived the concept. Chao Yu, Jun Qin, Xi Zhao, Shitong Luo, Hao Zeng, Bo Xu finished the experimental perform. Chao Yu and Jingjing Li generated the figures. Jingjing Li revised the manuscript. All authors reviewed and approved the manuscript.

## Data and materials availability

All data are available in the main text or the supplementary materials.

## Provenance and peer review

Not commissioned, externally peer-reviewed.

Supplementary Figure 1: A. 20 sample data before normalized. B. Overview of DEGs. C. Volcano map of DEGs. D. Variation trend map of DEGs. E. Display of the relationship between the main enriched KEGG pathway. F. Hub genes of DEGs (P<0.05, log2 ≥ 1 or ≤ - 1) were screened by Cytoscape.

## References

1. Hunter DJ, Schofield D, Callander E. The individual and socioeconomic impact of osteoarthritis. Nat Rev Rheumatol. 2014;10(7):437–441.

2. Martel-Pelletier J, Barr AJ, Cicuttini FM, et al. Osteoarthritis. Nat Rev Dis Primers. 2016;2:16072.

3. Loeser RF, Collins JA, Diekman BO. Ageing and the pathogenesis of osteoarthritis. Nat Rev Rheumatol. 2016;12(7):412–420.

4. Price AJ, Alvand A, Troelsen A, et al. Knee replacement. Lancet. 2018;392(10158):1672–1682.

5. Mastbergen SC, Ooms A, Turmezei TD, et al. Subchondral bone changes after joint distraction treatment for end stage knee osteoarthritis. Osteoarthritis Cartilage. 2022;30(7):965–972.

6. Sun Q, Zhen G, Li TP, et al. Parathyroid hormone attenuates osteoarthritis pain by remodeling subchondral bone in mice. Elife. 2021;10.

7. Cui Z, Wu H, Xiao Y, et al. Endothelial PDGF-BB/PDGFR-beta signaling promotes osteoarthritis by enhancing angiogenesis-dependent abnormal subchondral bone formation. Bone Res. 2022;10(1):58.

8. Hu W, Chen Y, Dou C, Dong S. Microenvironment in subchondral bone: predominant regulator for the treatment of osteoarthritis. Ann Rheum Dis. 2021;80(4):413–422.

9. Cai A, Hutchison E, Hudson J, et al. Metabolic enrichment of omega-3 polyunsaturated fatty acids does not reduce the onset of idiopathic knee osteoarthritis in mice. Osteoarthritis Cartilage. 2014;22(9):1301–1309.

10. Ross-Jones TN, McIlwraith CW, Kisiday JD, Hess TM, Hansen DK, Black J. Influence of an n-3 long-chain polyunsaturated fatty acid-enriched diet on experimentally induced synovitis in horses. J Anim Physiol Anim Nutr (Berl). 2016;100(3):565–577.

11. Lippiello L, Walsh T, Fienhold M. The association of lipid abnormalities with tissue pathology in human osteoarthritic articular cartilage. Metabolism. 1991;40(6):571–576.

12. Cillero-Pastor B, Eijkel G, Kiss A, Blanco FJ, Heeren RM. Time-of-flight secondary ion mass spectrometry-based molecular distribution distinguishing healthy and osteoarthritic human cartilage. Anal Chem. 2012;84(21):8909–8916.

13. Loef M, Schoones JW, Kloppenburg M, Ioan-Facsinay A. Fatty acids and osteoarthritis: different types, different effects. Joint Bone Spine. 2019;86(4):451–458.

14. McNulty AL, Miller MR, O’Connor SK, Guilak F. The effects of adipokines on cartilage and meniscus catabolism. Connect Tissue Res. 2011;52(6):523–533.

15. Jones JG. Hepatic glucose and lipid metabolism. Diabetologia. 2016;59(6):1098–1103.

16. Kastaniotis AJ, Autio KJ, Keratar JM, et al. Mitochondrial fatty acid synthesis, fatty acids and mitochondrial physiology. Biochim Biophys Acta Mol Cell Biol Lipids. 2017;1862(1):39–48.

17. Hara T, Kimura I, Inoue D, Ichimura A, Hirasawa A. Free fatty acid receptors and their role in regulation of energy metabolism. Rev Physiol Biochem Pharmacol. 2013;164:77–116.

18. Monfoulet LE, Philippe C, Mercier S, Coxam V, Wittrant Y. Deficiency of G-protein coupled receptor 40, a lipid-activated receptor, heightens in vitro- and in vivo-induced murine osteoarthritis. Exp Biol Med (Maywood). 2015;240(7):854–866.

19. Chen Y, Zhang D, Ho KW, et al. GPR120 is an important inflammatory regulator in the development of osteoarthritis. Arthritis Res Ther. 2018;20(1):163.

20. Sillat T, Barreto G, Clarijs P, et al. Toll-like receptors in human chondrocytes and osteoarthritic cartilage. Acta Orthop. 2013;84(6):585–592.

21. Bordji K, Grillasca JP, Gouze JN, et al. Evidence for the presence of peroxisome proliferator-activated receptor (PPAR) alpha and gamma and retinoid Z receptor in cartilage. PPARgamma activation modulates the effects of interleukin-1beta on rat chondrocytes. J Biol Chem. 2000;275(16):12243–12250.

22. Clockaerts S, Bastiaansen-Jenniskens YM, Feijt C, et al. Peroxisome proliferator activated receptor alpha activation decreases inflammatory and destructive responses in osteoarthritic cartilage. Osteoarthritis Cartilage. 2011;19(7):895–902.

23. Wachsmuth L, Soder S, Fan Z, Finger F, Aigner T. Immunolocalization of matrix proteins in different human cartilage subtypes. Histol Histopathol. 2006;21(5):477–485.

24. Turlip R, Sussman JH, Gujral J, et al. Toward transparency: Implications and future directions of artificial intelligence prediction model reporting in healthcare. Surg Neurol Int. 2025;16:135.

25. Williams GA, Callon KE, Watson M, et al. Skeletal phenotype of the leptin receptor-deficient db/db mouse. J Bone Miner Res. 2011;26(8):1698–1709.

26. Heep H, Wedemeyer C, Wegner A, Hofmeister S, von Knoch M. Differences in trabecular bone of leptin-deficient ob/ob mice in response to biomechanical loading. Int J Biol Sci. 2008;4(3):169–175.

27. Donell S. Subchondral bone remodelling in osteoarthritis. EFORT Open Rev. 2019;4(6):221–229.

28. Zhu X, Chan YT, Yung PSH, Tuan RS, Jiang Y. Subchondral Bone Remodeling: A Therapeutic Target for Osteoarthritis. Front Cell Dev Biol. 2020;8:607764.

29. Sun HB. Mechanical loading, cartilage degradation, and arthritis. Ann N Y Acad Sci. 2010;1211:37–50.

30. Medvedeva EV, Grebenik EA, Gornostaeva SN, et al. Repair of Damaged Articular Cartilage: Current Approaches and Future Directions. Int J Mol Sci. 2018;19(8).

31. Chen X, Du J, Zhan W, et al. Polyene phosphatidylcholine promotes tibial fracture healing in rats by stimulating angiogenesis dominated by the VEGFA/VEGFR2 signaling pathway. Biochem Biophys Res Commun. 2024;719:150100.

32. Wei G, Lu K, Umar M, et al. Risk of metabolic abnormalities in osteoarthritis: a new perspective to understand its pathological mechanisms. Bone Res. 2023;11(1):63.

33. Gkretsi V, Simopoulou T, Tsezou A. Lipid metabolism and osteoarthritis: lessons from atherosclerosis. Prog Lipid Res. 2011;50(2):133–140.

34. Masuko K, Murata M, Suematsu N, et al. A metabolic aspect of osteoarthritis: lipid as a possible contributor to the pathogenesis of cartilage degradation. Clin Exp Rheumatol. 2009;27(2):347–353.

35. Aziz A, Ganesan Nathan K, Kamarul T, Mobasheri A, Sharifi A. The interplay between dysregulated metabolites and signaling pathway alterations involved in osteoarthritis: a systematic review. Ther Adv Musculoskelet Dis. 2024;16:1759720X241299535.

36. Adam MS, Zhuang H, Ren X, Zhang Y, Zhou P. The metabolic characteristics and changes of chondrocytes in vivo and in vitro in osteoarthritis. Front Endocrinol (Lausanne). 2024;15:1393550.

37. Wang Y, Wei L, Zeng L, He D, Wei X. Nutrition and degeneration of articular cartilage. Knee Surg Sports Traumatol Arthrosc. 2013;21(8):1751–1762.

38. Santos AL, Preta G. Lipids in the cell: organisation regulates function. Cell Mol Life Sci. 2018;75(11):1909–1927.

39. Arkill KP, Winlove CP. Fatty acid transport in articular cartilage. Arch Biochem Biophys. 2006;456(1):71–78.

40. Shin Y, Huh YH, Kim K, et al. Low-density lipoprotein receptor-related protein 5 governs Wnt-mediated osteoarthritic cartilage destruction. Arthritis Res Ther. 2014;16(1):R37.

41. Toussirot E, Streit G, Wendling D. The contribution of adipose tissue and adipokines to inflammation in joint diseases. Curr Med Chem. 2007;14(10):1095–1100.

42. Binvignat M, Sellam J, Berenbaum F, Felson DT. The role of obesity and adipose tissue dysfunction in osteoarthritis pain. Nat Rev Rheumatol. 2024;20(9):565–584.

43. Saponaro C, Gaggini M, Carli F, Gastaldelli A. The Subtle Balance between Lipolysis and Lipogenesis: A Critical Point in Metabolic Homeostasis. Nutrients. 2015;7(11):9453–9474.

44. Zhu M, Tang D, Wu Q, et al. Activation of beta-catenin signaling in articular chondrocytes leads to osteoarthritis-like phenotype in adult beta-catenin conditional activation mice. J Bone Miner Res. 2009;24(1):12–21.

45. Giaginis C, Giagini A, Theocharis S. Peroxisome proliferator-activated receptor-gamma (PPAR-gamma) ligands as potential therapeutic agents to treat arthritis. Pharmacol Res. 2009;60(3):160–169.

46. Chen YJ, Chan DC, Lan KC, et al. PPARgamma is involved in the hyperglycemia-induced inflammatory responses and collagen degradation in human chondrocytes and diabetic mouse cartilages. J Orthop Res. 2015;33(3):373–381.

47. Monemdjou R, Vasheghani F, Fahmi H, et al. Association of cartilage-specific deletion of peroxisome proliferator-activated receptor gamma with abnormal endochondral ossification and impaired cartilage growth and development in a murine model. Arthritis Rheum. 2012;64(5):1551–1561.

48. Liu Q, Li M, Wang S, Xiao Z, Xiong Y, Wang G. Recent Advances of Osterix Transcription Factor in Osteoblast Differentiation and Bone Formation. Front Cell Dev Biol. 2020;8:601224.

49. Amarasekara DS, Kim S, Rho J. Regulation of Osteoblast Differentiation by Cytokine Networks. Int J Mol Sci. 2021;22(6).

50. Liu X, Zhang P, Gu Y, Guo Q, Liu Y. Type H vessels: functions in bone development and diseases. Front Cell Dev Biol. 2023;11:1236545.

51. Li YJ, Guo Q, Ye MS, et al. YBX1 promotes type H vessel-dependent bone formation in an m5C-dependent manner. JCI Insight. 2024;9(4).

52. Vasheghani F, Zhang Y, Li YH, et al. PPARgamma deficiency results in severe, accelerated osteoarthritis associated with aberrant mTOR signalling in the articular cartilage. Ann Rheum Dis. 2015;74(3):569–578.

53. Fan J, Xie Y, Liu D, et al. Crosstalk Between H-Type Vascular Endothelial Cells and Macrophages: A Potential Regulator of Bone Homeostasis. J Inflamm Res. 2025;18:2743–2765.

54. He Y, Bundkirchen K, Taheri S, et al. Increased vascularization of the subchondral region in human osteoarthritic femoral head in the elderly. Histochem Cell Biol. 2025;163(1):39.

55. Olah T, Cucchiarini M, Madry H. Temporal progression of subchondral bone alterations in OA models involving induction of compromised meniscus integrity in mice and rats: A scoping review. Osteoarthritis Cartilage. 2024;32(10):1220–1234.

56. Zou Y, Wang Z, Shi H, Hu J, Hu W. Soybean Isoflavones Alleviate Osteoarthritis Through Modulation of the TSC1/mTORC1 Signaling Pathway to Reduce Intrachondral Angiogenesis. Immunol Invest. 2024;53(8):1439–1455.

57. Liu Y, Xie HQ, Shen B. Type H vessels-a bridge connecting subchondral bone remodelling and articular cartilage degeneration in osteoarthritis development. Rheumatology (Oxford). 2023;62(4):1436–1444.

58. Peng Y, Wu S, Li Y, Crane JL. Type H blood vessels in bone modeling and remodeling. Theranostics. 2020;10(1):426–436.

59. Heidari B. Knee osteoarthritis prevalence, risk factors, pathogenesis and features: Part I. Caspian J Intern Med. 2011;2(2):205–212.

60. Kilkenny C, Browne WJ, Cuthill IC, Emerson M, Altman DG. Improving bioscience research reporting: The ARRIVE guidelines for reporting animal research. J Pharmacol Pharmacother. 2010;1(2):94–99.

